# Vtc5 is localized to the vacuole membrane by the conserved AP-3 complex to regulate polyphosphate synthesis in budding yeast

**DOI:** 10.1101/2021.03.15.434853

**Authors:** Amanda Bentley-DeSousa, Michael Downey

## Abstract

Polyphosphates (polyP) are energy-rich polymers of inorganic phosphates assembled into chains ranging from 3-1000s of residues in length. They are thought to exist in all cells on earth and play roles in an eclectic mix of functions ranging from phosphate homeostasis to cell signaling, infection control, and blood clotting. In the budding yeast *Saccharomyces cerevisiae*, polyP chains are synthesized by the vacuole-bound VTC complex, which synthesizes polyP while simultaneously translocating it into the vacuole lumen where it is stored at high concentrations. VTC’s activity is promoted by an accessory subunit called Vtc5. In this work, we find that the conserved AP-3 complex is required for proper Vtc5 localization to the vacuole membrane. In human cells, previous work has demonstrated that mutation of AP-3 subunits gives rise to Hermansky-Pudlak Syndrome, a rare disease with molecular phenotypes that include decreased polyP accumulation in platelet dense granules. In yeast AP-3 mutants, we find that Vtc5 is rerouted to the vacuole lumen by the ESCRT complex, where it is degraded by the vacuolar protease Pep4. Cells lacking functional AP-3 have decreased levels of polyP, demonstrating that membrane localization of Vtc5 is required for its VTC stimulatory activity *in vivo*. Our work provides insight into the molecular trafficking of a critical regulator of polyP metabolism in yeast. We speculate that AP-3 may also be responsible for the delivery of polyP regulatory proteins to platelet dense granules in higher eukaryotes.

**HIGHLIGHTS:** Vtc5 localization to the vacuole membrane depends on the AP-3 complex

The ESCRT pathway brings mislocalized Vtc5 to the vacuole lumen where it is degraded

Decreased polyP levels in AP-3 mutants are explained by Vtc5 mislocalization

Deletion of *DOA4* restores wild-type localization of Vtc5 without restoring polyP levels

## INTRODUCTION

Polyphosphates (polyP) are chains of inorganic phosphates found in all cell types studied to date. PolyP chains are variable in size, ranging from 3-1000s of units in length, and are linked together via high-energy phosphoanhydride bonds^1^. While once dismissed as ‘molecular fossils’, recent work suggests that polyP plays critical roles in diverse processes across both prokaryotic and eukaryotic organisms including bacterial virulence, infection control, blood coagulation, protein folding, and diverse aspects of cell signaling^2-7^. As such, polyP chains have gained significant interest as a potential target for therapeutics in a wide variety of pathologies. In contrast to bacteria and fungi, the enzymes that synthesize polyP chains in higher eukaryotic cells are largely unknown. There are no clear homologs of either prokaryotic or fungal polyP synthetases in mammals. Recent work by the Abramov group suggests that the mammalian mitochondrial F_0_F_1_ ATPase has polyP synthesis capabilities^8^, but the overall contribution of this enzyme to total cellular pools of polyP remains to be tested.

One model organism used to study polyP at a foundational level is the budding yeast *S. cerevisiae*. Here, polyP is present in high concentrations (>200 mM), and the enzymes responsible for its metabolism have been identified^9-11^. In yeast, polyP is synthesized by the vacuole transporter chaperone (VTC) complex. The minimal (core) VTC complex is composed of 3 subunits: Vtc1, Vtc2 or Vtc3, and Vtc4 (the catalytic enzyme)^9, 12^. The Vtc3 subunit mostly localizes to the vacuole membrane alongside Vtc1 and Vtc4^12^. On the other hand, the Vtc2 subunit is found at the endoplasmic reticulum and/or cell periphery, and relocalizes to the vacuole membrane under phosphate-limited conditions^12^. PolyP synthesis by VTC requires simultaneous translocation of polyP into the vacuole lumen^13^, where it makes up over 10 % of the dry weight of the cell^10^. PolyP is also found in lower concentrations in the cytoplasm, plasma membrane, mitochondria, and nucleus, although observed concentrations vary by method of analysis^14^. In addition to being the only known polyP synthetase in yeast, the VTC complex has been demonstrated or suggested to play a role in a myriad of cellular activities such as the stability of vacuolar V-ATPase subunits, in vacuole fusion, and in microautophagy^15-17^. PolyP itself has also been implicated in the regulation of pH balance^18^, ion homeostasis^19-21^, and phosphate metabolism^18, 22-24^.

Recent work has identified several exciting aspects of VTC regulation. First, activity of the complex is promoted by binding of inositol pyrophosphate InsP7 to the SPX domains of VTC proteins^25, 26^. As such, loss of Kcs1, which catalyzes the formation of InsP7, drastically reduces polyP levels^27, 28^. Although canonical (serine/threonine) phosphorylation and lysine ubiquitylation sites have been identified on multiple VTC subunits^29-31^, their functions are currently unknown. Recently, a new VTC regulatory subunit termed Vtc5 was identified^22^. Vtc5 localizes exclusively to the vacuole membrane and interacts with the VTC complex to increase the rate of polyP production^22^. PolyP levels in *vtc5*Δ mutants are reduced to 20% of those of wild-type cells^22^. The mechanism by which Vtc5 exerts its positive effects on VTC is unknown, although it appears to function independently of effects imparted by inositol pyrophosphates^22^. Given the importance of Vtc5 as a regulator of the core VTC complex, we sought to identify the pathways that are responsible for localizing Vtc5 to the vacuole membrane.

Membrane proteins are synthesized by ribosomes at the endoplasmic reticulum and transported to the trans-golgi network (TGN) prior to being sorted to their final destination^32^. Vacuolar proteins are sorted by two well established protein transport pathways, the CPY (indirect / <u>C</u>arboxy<u>p</u>eptidase <u>Y</u>) or AP-3 (direct / <u>A</u>daptor <u>P</u>rotein Complex <u>3</u>) transport pathways. The CPY pathway transports cargoes to the vacuole in an indirect fashion using the endosomal system as an intermediate path. In this pathway, cargoes localize to endosomes prior to fusion with the vacuole membrane for delivery (**Fig. 1A**)^32^. In contrast, AP-3 cargoes are selected at the TGN, where the complex buds from the TGN with its cargoes to create AP-3 coated vesicles. These vesicles are transported directly to the vacuole through the cytoplasm, and AP-3 docks at the vacuole membrane to release its cargoes (**Fig. 1A**)^32^. There are links between the AP-3 complex and polyP storage in higher eukaryotes. In humans, mutations in AP-3 are associated with a rare disorder called Hermansky-Pudlak Syndrome^33-35^. Hermansky-Pudlak Syndrome patients lack the ability to synthesize lysosome-related organelles in diverse cell types throughout the body, which results in broad phenotypes that include albinism, visual impairment, and bleeding problems^35^. Notably, this includes the inability to generate dense granules in platelets, where polyP is usually stored in its highest concentrations (130 mM)^36^. PolyP promotes blood clotting by acting on proteins involved in the coagulation cascade including factor XII^37^, factor XI/V^38, 39^, and thrombin^40^. PolyP addition re-establishes clotting stimulatory activity of platelets derived from Hermansky-Pudlak patients^41^.

**Figure 1:**
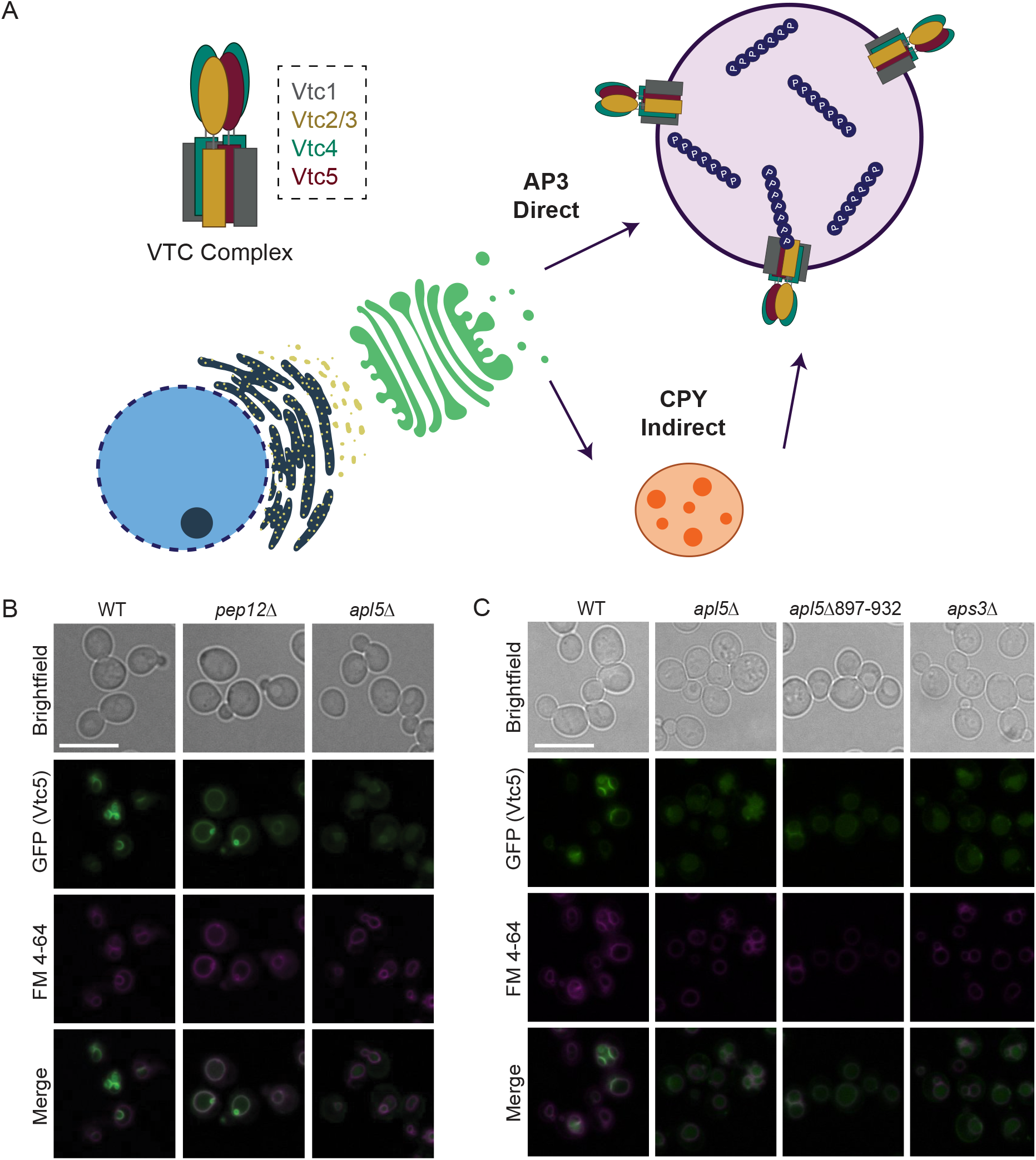
GFP-Vtc5 is localized to the vacuole membrane via the conserved AP-3 pathway. **A**) Simplified schematic of protein transport to the vacuole in *S. cerevisiae*. **B**) The indicated strains were grown in YPD prior to incubation with FM 4-64, which marks the vacuole membrane, for 2 hours. Cells were then washed with fresh YPD for 30 minutes, transferred to synthetic media, and imaged. Live-cell fluorescence microscopy was performed using a Leica DMI 6000 microscope at 63X with oil immersion. For AP-3 mutants, green (GFP-Vtc5) images were taken at longer exposure times to account for differences in signal intensity. Images were processed in FIJI. Scale bar represents 10 μm. **C**) The indicated strains were processed as in B.

In this study, we describe a role for the yeast AP-3 complex in localizing Vtc5 to the vacuole membrane. In AP-3 mutants, Vtc5 is rerouted to the vacuole lumen by the <u>E</u>ndosomal <u>S</u>orting <u>C</u>omplex <u>R</u>equired for <u>T</u>ransport (ESCRT) complex where it is degraded by the vacuolar protease Pep4. Mislocalization of Vtc5 in AP-3 mutants is accompanied by a decrease in VTC protein levels and decreased levels of polyP. Overall, our study explains the polyP accumulation defects in AP-3 mutants and provides novel insights into the regulation of the VTC polyP synthetase in budding yeast.

## RESULTS

### Vtc5 localization to the vacuole membrane is disrupted in AP-3 mutants

The CPY and AP-3 pathways comprise the two major routes of protein transport to the vacuole from the TGN (**Fig. 1A**). To test which of these pathways is responsible for Vtc5 localization to the vacuole membrane, we used live-cell fluorescence microscopy to analyze GFP-Vtc5 localization in AP-3 and CPY pathway mutants. Notably, all N-terminal GFP-Vtc fusions used in this work are functional, with the GFP tag facing the cytoplasmic side of the vacuole membrane to ensure the tag is not degraded within the vacuole lumen^12, 22^. These GFP fusions are expressed from constitutive promoters integrated at endogenous *VTC* loci (See Experimental Procedures). In wild-type cells, GFP-Vtc5 (green) localized exclusively to the vacuole membrane as demonstrated by colocalization with FM4-64 (magenta), a dye which labels the vacuole membrane (**Fig. 1B**). CPY (*pep12*Δ) mutants showed GFP-Vtc5 localization similar to that of wild-type cells (**Fig. 1B**). Conversely, GFP-Vtc5 was mislocalized to the vacuole lumen in AP-3 mutant (*apl5*Δ) cells (**Fig 1B**). AP-3 is highly conserved from yeast to humans. It consists of a heterotetramer of four protein subunits: two large subunits (Apl5/AP3β1and Apl6/AP3δ1), one medium subunit (Apm3/AP3μ1), and one small subunit (Aps3/AP3σ1) (**Supplemental Fig. 1A**)^42^. Deletion of genes encoding any one of the four AP-3 subunits results in defects in AP-3 cargo transport^43^. Therefore, to corroborate our results with *apl5*Δ, we examined GFP-Vtc5 localization in *aps3*Δ cells and again observed its mislocalization to the vacuole lumen in this mutant (**Fig. 1C**). Overall, these data suggest that the primary pathway responsible for localizing GFP-Vtc5 to the vacuole membrane is the AP-3 transport pathway, with little if any compensation by the CPY pathway.

Intriguingly, polyP chains can be covalently attached to protein targets as a post-translational modification termed polyphosphorylation^27, 44^. Polyphosphorylation is the non-enzymatic addition of polyP chains onto lysine residues, principally within poly-acidic, serine, and lysine (PASK) rich clusters^27, 45^. We previously reported that Apl5 is polyphosphorylated in its C-terminus PASK cluster (amino acids 897-932)^46^. Therefore, to test if polyphosphorylation impacts Apl5’s role in localizing GFP-Vtc5 to the vacuole membrane, we first deleted the PASK cluster in its entirety. Deletion of Apl5’s PASK cluster (*apl5*Δ897-932) resulted in mislocalization of GFP-Vtc5 to the vacuole lumen, similar to what we observed in AP-3 null mutants (**Fig. 1C**). To define the contribution of polyphosphorylation more specifically, we generated a strain wherein Apl5 is expressed with 13 lysine to arginine (K-R) amino acid substitutions in its PASK cluster. Polyphosphorylation results in an electrophoretic shift of target proteins separated on Bis-Tris NuPAGE gels^27, 47^, and this is currently the only method described for evaluation of this new modification. As demonstrated by NuPAGE analysis, the resulting mutant (Apl5_PASK_K-R) was unable to undergo polyphosphorylation. (**Supplemental Fig. 1B**).

However, this did not impact GFP-Vtc5 localization (**Supplemental Fig. 1C**), nor did it mimic other known *apl5*Δ phenotypes such as defects in the maturation of AP-3 target vacuolar alkaline phosphatase Pho8^48^ (**Supplemental Fig. 1D**), or enhanced sensitivity to nickel chloride^49^ or rapamycin^50^ (**Supplemental Fig. 2A-B**). We note that the PASK cluster lies within Apl5’s Vps41 binding domain. Vps41 is a member of the H<u>o</u>motypic <u>F</u>usion and <u>P</u>rotein <u>S</u>orting (HOPS) complex required for vesicle docking at the vacuole and delivery of AP-3 cargoes^51, 52^. As such, the role of the Apl5 PASK in GFP-Vtc5 delivery may stem from disruption of this interaction rather than a defect in polyphosphorylation. Interestingly, the Apl5 C-terminal PASK deletion mutant (*apl5*Δ897-932) showed defects in maturation of Pho8, but it did not result in sensitivity to nickel chloride or rapamycin (**Supplemental Figs. 1D, 2A-B**). This separation of function mutant may serve as a useful tool to dissect AP-3’s role underlying these distinct phenotypes.

**Figure 2:**
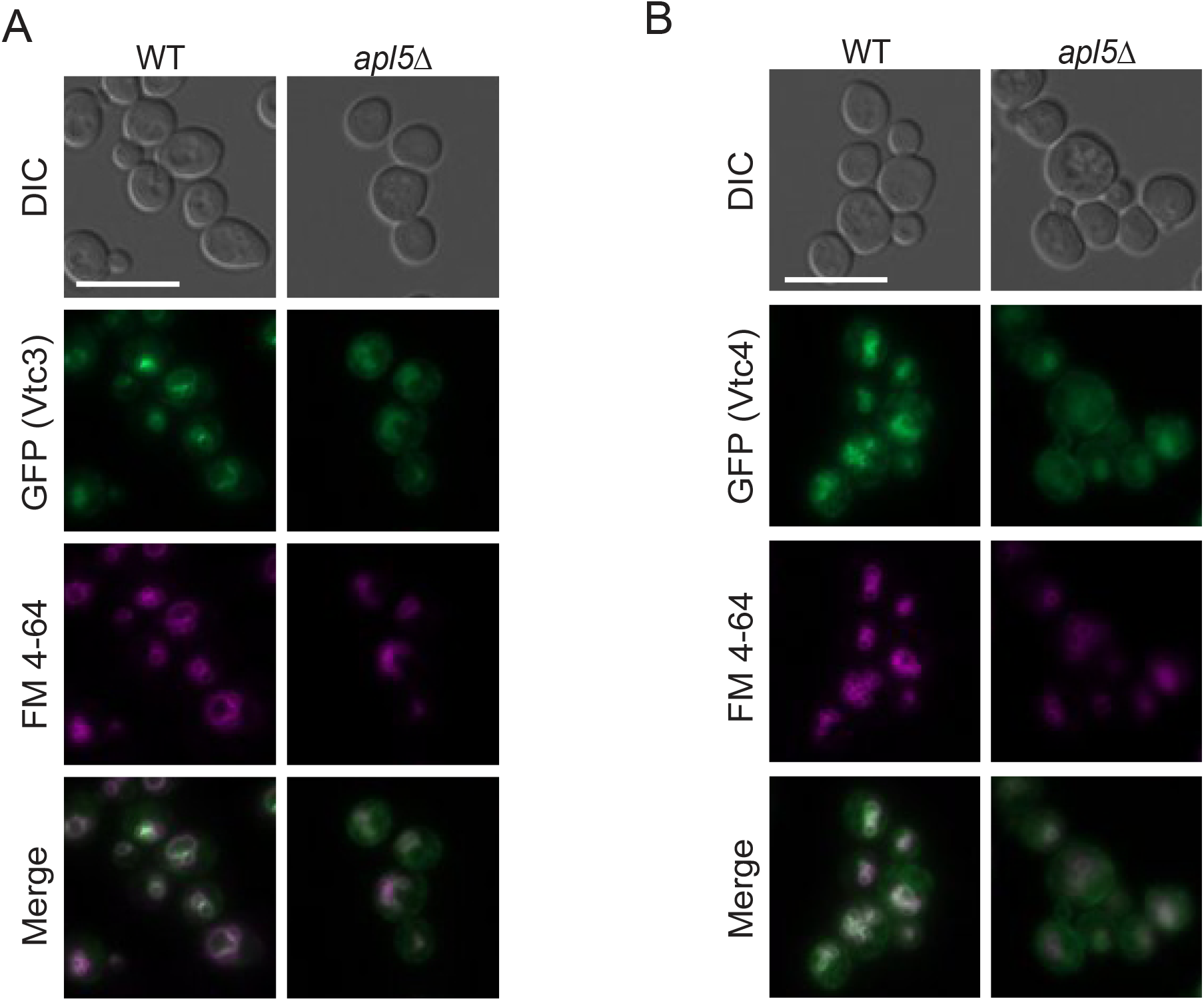
Partial disruption of GFP-Vtc3 and GFP-Vtc4 localization in AP-3 mutants. **A)** The indicated strains were grown in YPD media prior to incubation with FM 4-64, which marks the vacuole membrane, for 2 hours. Cells were then washed with fresh YPD for 30 minutes, transferred to synthetic media, and imaged. Live-cell fluorescence microscopy was performed using a Zeiss AxioObserver 7 at 63X with oil immersion. Images were processed in FIJI. **B)** The indicated strains were processed as in A. Scale bar represents 10 μm.

We next sought to test whether vacuolar localization of additional VTC subunits was also impacted by *APL5* deletion. We observed no difference in the localization of GFP-Vtc3 in wild-type versus *apl5*Δ mutants (**Fig. 2A**). GFP-Vtc4 showed an intermediate phenotype, with increased localization to the cytoplasm in *apl5*Δ, with some protein also remaining at the vacuole membrane (**Fig. 2B**). These data suggest that although the AP-3 pathway is responsible for the proper vacuolar localization of GFP-Vtc5, other pathways may contribute to the localization of core VTC subunits. Given the importance of Vtc5 as a regulator of VTC activity and the clear disruption of its localization in AP-3 mutants, we focused on the transport of this subunit.

### GFP-Vtc subunits are degraded in cells lacking functional AP-3

We next used western blotting to gain insight into the fate of mislocalized GFP-Vtc5. Relative to wild-type controls, *apl5*Δ and *aps3*Δ cells showed a striking accumulation of free GFP, which is known to be resistant to degradation (**Fig. 3A-B**)^53^. This was not the result of increased protein expression, as full length GFP-Vtc5 was reduced in these mutants (**Fig. 3A & 3C**). This same pattern has been observed previously for other proteins mislocalized to the vacuole lumen and is attributed to cargo degradation^54, 55^. We also tested if GFP-Vtc4 and GFP-Vtc3 levels were similarly affected. Both GFP-Vtc4 and GFP-Vtc3 also accumulated free GFP at the expense of decreased full-length fusions, although the effect was not as dramatic as that observed for GFP-Vtc5 (**Fig. 3D-3I**). Previous work from the Mayer group showed that *vtc5*Δ cells have decreased levels of core VTC subunits at the vacuole membrane^22^. Degradation of GFP-Vtc3 and GFP-Vtc4, and partial mislocalization of GFP-Vtc4 in *apl5*Δ cells, are consistent with a model wherein mislocalized Vtc5 is largely non-functional in the absence of AP-3. However, we cannot exclude that these molecular phenotypes stem in part from defects in the transport of other AP-3 cargos (see Discussion).

**Figure 3:**
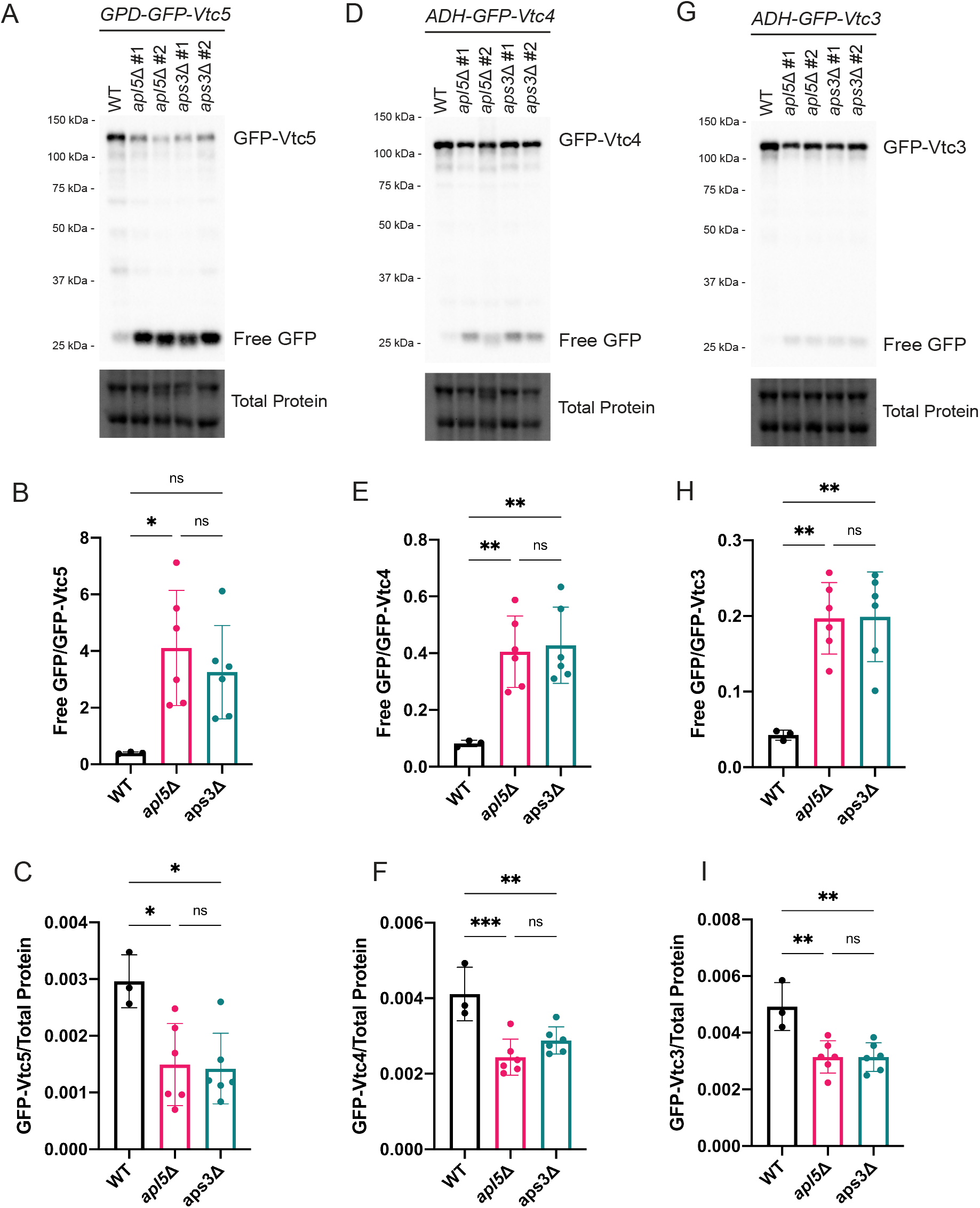
AP-3 mutation causes degradation of GFP-Vtc proteins. **A**) Upon AP-3 mutation (*apl5*Δ and *aps3*Δ), full-length GFP-Vtc5 protein levels are reduced with a concomitant increase in free GFP. Proteins were extracted from the indicated strains using a TCA protein extraction protocol, separated on a 10% BioRad TGX Stain-Free™ FastCast™ acrylamide gel, and transferred to a nitrocellulose membrane. The membrane was imaged on a BioRad ChemiDoc after immunoblotting with anti-GFP. Total protein was imaged as a loading control. **B-C)** Quantification of free GFP/GFP-Vtc5 ratios and GFP-Vtc5 full length levels. Quantifications were done using BioRad ImageLab software and graphs were created using Prism GraphPad Software. One-way ANOVAs were performed with Tukey posthoc tests. * *p* < 0.05, ** *p* < 0.01, *** *p* <0.001. Error bars represent standard deviation of the mean. n = 3 for WT and n = 6 for AP-3 mutants. **D)-F)** In AP-3 mutants, GFP-Vtc4-expressing strains accumulate free GFP at the expense of full-length protein. Methodology was as in A-C. **G)-I)** In AP-3 mutants, GFP-Vtc3-expressing strains accumulate free GFP at the expense of full-length protein. The same methodology was used as in 4A-C.

### Mislocalized Vtc5 is rerouted to the vacuole lumen by the ESCRT pathway

We next investigated which pathways are responsible for localizing GFP-Vtc5 to the vacuole lumen in the absence of functional AP-3, with a focus on the autophagy and ESCRT pathways. Autophagy is a process whereby cytoplasmic material becomes sequestered in vesicles that fuse to the vacuole to deliver their contents for degradation^56^. The ESCRT pathway is responsible for detecting ubiquitylated transmembrane proteins to sort them into multi-vesicular bodies (MVBs) in the endocytic pathway for delivery to the vacuole^57^. Disruption of ESCRT (*vps27*Δ), but not autophagy (*atg8*Δ) in AP-3 mutants resulted in a loss of GFP signal from the vacuole lumen (**Fig. 4A**). In contrast, neither pathway impacted GFP-Vtc5 localization in wild-type cells (**Supplemental Fig. 3A**). When ESCRT complex subunits are mutated, the complex becomes defective for proper MVB biogenesis and fusion at the vacuole membrane^57^. Indeed, in *apl5*Δ *vps27*Δ double mutants, GFP-Vtc5 appeared to accumulate in FM4-64 labelled MVBs around the vacuole membrane (**Fig. 4A**). Deletion of *VPS27* also resulted in reversal of free GFP accumulation in western blots (**Fig. 4B**). This same molecular phenotype was observed across multiple ESCRT complex mutants (ESCRT-0, -I, -II, and -III, **Supplemental Fig. 3B)**.

**Figure 4:**
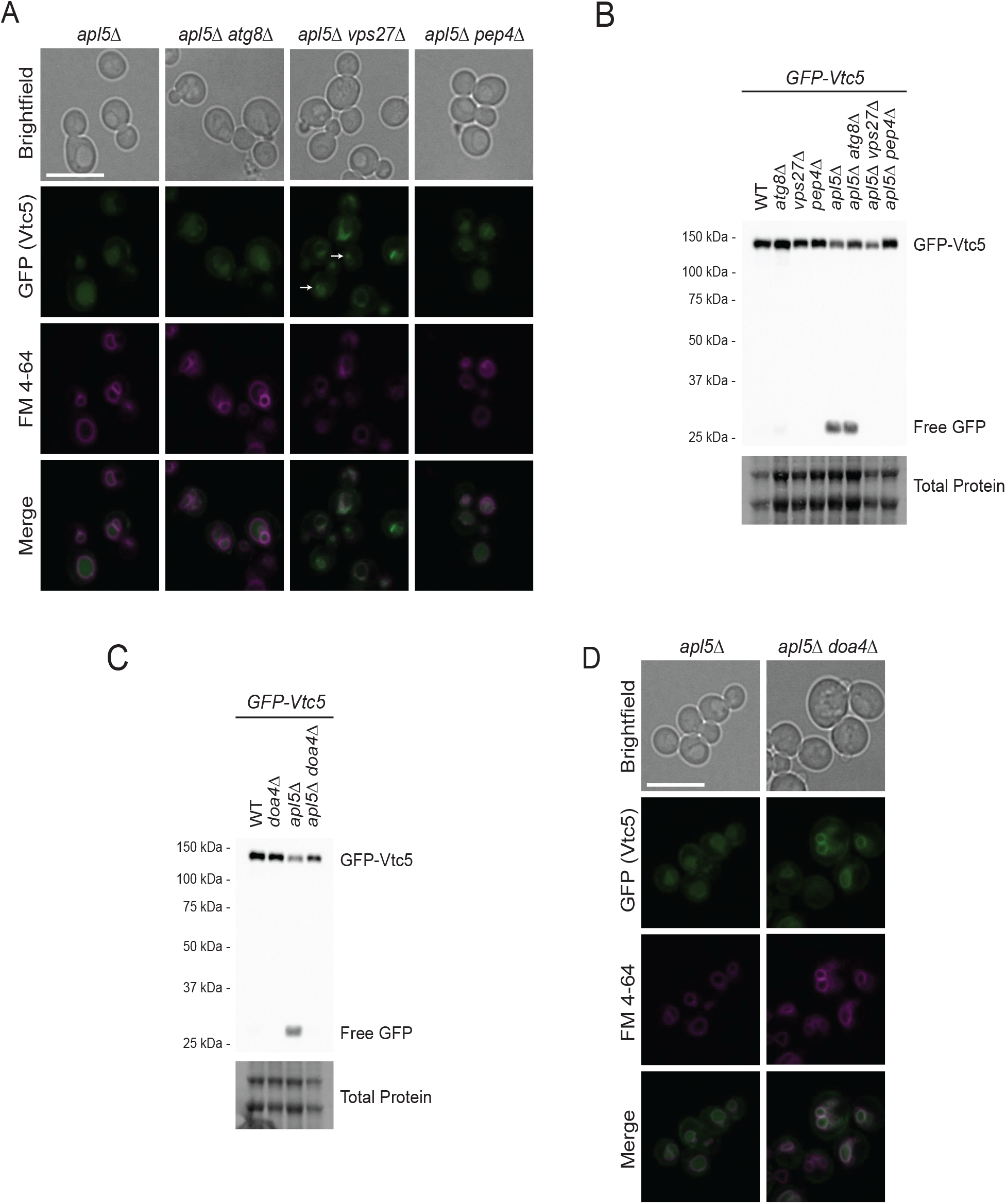
Degradation of mislocalized GFP-Vtc5 depends on ESCRT and the Pep4 protease. **A)** The mislocalization of GFP-Vtc5 upon AP-3 mutation is mediated by the ESCRT pathway. Cells were grown in YPD prior to incubation with FM4-64, which marks the vacuole membrane, for 2 hours. Cells were then washed with fresh YPD for 30 minutes, transferred to synthetic media, and imaged. Live-cell fluorescence microscopy was performed using a Leica DMI 6000 at 63X with oil immersion. Images were processed in FIJI. Scale bar represents 10 μm. White arrows indicate potential MVBs at vacuole. **B)** Free GFP accumulation in GFP-Vtc5, mediated by AP-3 mutation, is reversed by ESCRT (*vps27*Δ) and *pep4*Δ mutation. Proteins from the indicated strains were extracted using a TCA protein extraction protocol, separated on a 10% BioRad TGX Stain-Free™ FastCast™ acrylamide gel, and transferred to a nitrocellulose membrane. The membrane was imaged for total protein and probed using an anti-GFP antibody to detect GFP-Vtc5 protein using a BioRad ChemiDoc. **C)** GFP-Vtc5 degradation and free GFP accumulation is reversed by *DOA4* deletion. The same methodology was used as in 4B. **D)** Deletion of *DOA4* rescues localization of GFP-Vtc5 in *apl5*Δ mutants. The same methodology was used as in 4A.

The ESCRT pathway recognizes cargoes that have been ubiquitylated by E3 ubiquitin ligases^57, 58^. These cargoes are then transported to the vacuole via MVBs that fuse with the vacuole membrane to deliver its contents. As one of the last steps in this process, the deubiquitinase Doa4 removes ubiquitin moieties from ESCRT-targeted proteins in order to recycle ubiquitin levels within the cell upon cargo delivery to the vacuole^57, 58^. Mutation of *DOA4* alone has no impact on GFP-Vtc5 processing or localization (**Fig. 4C and Supplemental Fig. 3C**). In AP-3 mutants, however, *doa4*Δ prevented the accumulation free GFP, mirroring what was seen for ESCRT mutants (**Fig. 4C**). However, in contrast to the ESCRT mutants, *doa4*Δ also restored robust GFP-Vtc5 localization to the vacuole membrane (**Fig. 4D**).

Altogether, we conclude that Doa4 is a key player in localizing Vtc5 to the vacuole lumen in the absence of AP-3. Finally, when proteins accumulate in the lumen, they become accessible to vacuole proteases^59^. Deletion of the major vacuole protease Pep4 largely rescued both full-length GFP-Vtc5 and reversed the accumulation of free GFP in AP-3 mutants (**Fig. 4A-B**), suggesting that Pep4 is a major protease responsible for Vtc5 degradation.

### The AP-3 complex is required for maintenance of wild-type polyP levels

Finally, we tested the contribution of AP-3 to polyP homeostasis. Deletion of AP-3 subunits resulted in a clear decrease in polyP levels (**Fig. 5A**). This finding is consistent with previous data wherein AP-3 mutants were found to be important for polyP accumulation in a large scale screen^60^. We observed a similar result in our strains used for prior analyses where GFP-Vtc5 is expressed under a constitutive *GPD1* promoter (**Fig. 5B**). Together with our previous results, these data suggest that correct localization of Vtc5 to the vacuole membrane by the AP-3 complex is important for its function. Notably, AP-3 mutants still have more polyP than cells lacking Vtc5 altogether (*vtc5*Δ, **Fig. 5C**). This observation could be explained by residual localization of Vtc5 to the vacuole membrane in AP-3 mutants. Interestingly, deletion of *DOA4*, which rescues GFP-Vtc5 protein levels and localization to the vacuole membrane, was unable to reverse the decreased polyP levels in *apl5*Δ mutants (**Fig. 5D**). Thus, proper function of GFP-Vtc5 requires localization to the vacuole membrane, specifically through the AP-3 pathway.

**Figure 5:**
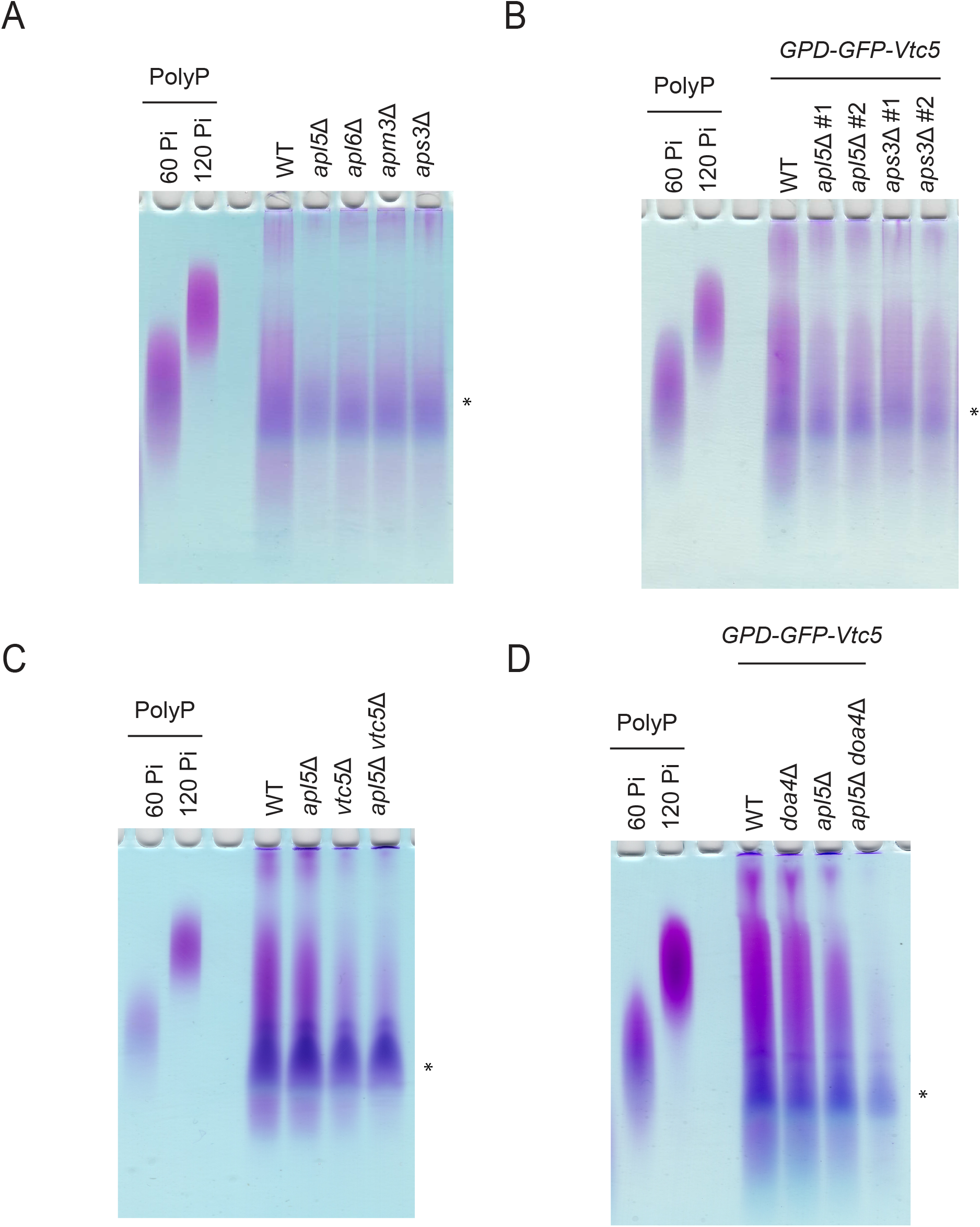
Functional AP-3 is required for the maintenance of polyP levels. **A)** AP-3 subunit mutant causes a reduction in polyP levels. There is a loss of polyP when mutating each of the four AP-3 subunits (*apl5*Δ, *apl6*Δ, *apm3*Δ, and *aps3*Δ). PolyP was extracted as described in Experimental Procedures. After extraction, samples were mixed with polyP sample buffer and separated on a 15.8% TBE-Urea Acrylamide gel. The gel was incubated in fixing solution with toluidine blue for 15 minutes prior to destaining and imaging. PolyP standards (14, 60, and 120 Pi units) are included as standards. **B)** AP-3 mutants (*apl5*Δ and *aps3*Δ) cause a reduction in polyP levels when GFP-Vtc5 is expressed under a *GPD1* promoter. The same methodology was used as in 5A. **C)** Mutating both AP-3 (*apl5*Δ) and *vtc5*Δ results in a largely epistatic decrease in polyP levels. The same methodology was used as in 5A. **D)** Although *doa4*Δ is capable of rescuing GFP-Vtc5 localization and protein degradation when AP-3 is mutated (*apl5*Δ), it is incapable of restoring polyP levels. The same methodology was used as in 5A. Asterisk indicates a background contaminant in extractions.

## DISCUSSION

### AP-3 regulation of Vtc5 localization

Interest in polyP research has experienced a resurgence in recent years on the heels of exciting connections between polyP and diverse aspects of cell signaling and protein homeostasis. With the key players involved in mammalian polyP metabolism largely uncharacterized, model systems have proved essential to our understanding of polyP dynamics and function. In *S. cerevisiae*, polyP is synthesized by the vacuole-bound VTC complex and stored at high concentrations in the vacuole^13^. The activity of the core VTC complex (consisting of Vtc1, Vtc2 or Vtc3, and Vtc4) is increased dramatically by the Vtc5 subunit^22^. This is the only protein known to act directly on the VTC complex to stimulate polyP production. Our study supports a model where Vtc5 localization to the vacuole membrane depends on the evolutionarily conserved AP-3 complex. This finding provides insight into the regulation of the VTC complex and identifies Vtc5 as a cargo whose mislocalization underlies polyP accumulation defects observed in AP-3 mutants.

In wild-type cells (**Fig. 6)**, we propose that Vtc5 is sorted into AP-3 coated vesicles at the TGN and transported directly to the vacuole. The AP-3 complex selects protein cargoes by tyrosine-based motifs (YXXØ where X represents any amino acid and Ø is a bulky hydrophobic amino acid) and/or dileucine-based motifs ([D/E]XXXL[L/I] where X represents any amino acid)^61^. Notably, Vtc5 has 6 tyrosine-based motifs and 2 dileucine-based motifs which together could be used as a targeting signal. Alternatively, Vtc5 could be transported in conjunction with other AP-3 cargoes such as Vam3, which is also required for wild-type levels of polyP accumulation^60^. Regardless, we propose that AP-3 coated vesicles may bind to Vps41 of the HOPS complex to facilitate docking and release of cargo, including Vtc5, into the vacuole membrane. In the absence of AP-3, Vtc5 is mislocalized to the vacuole lumen. Mislocalized Vtc5 (e.g. in *apl5*Δ), is recognized and sorted by the ESCRT pathway into the vacuole lumen via the endosomal system where it is eventually degraded in a manner dependent on vacuole protease Pep4. Since Pep4 is also required for the activation of vacuole proteases Prb1 and Prc1^62, 63^, it is possible that these also these play a role in Vtc5 degradation in the vacuole lumen.

**Figure 6:**
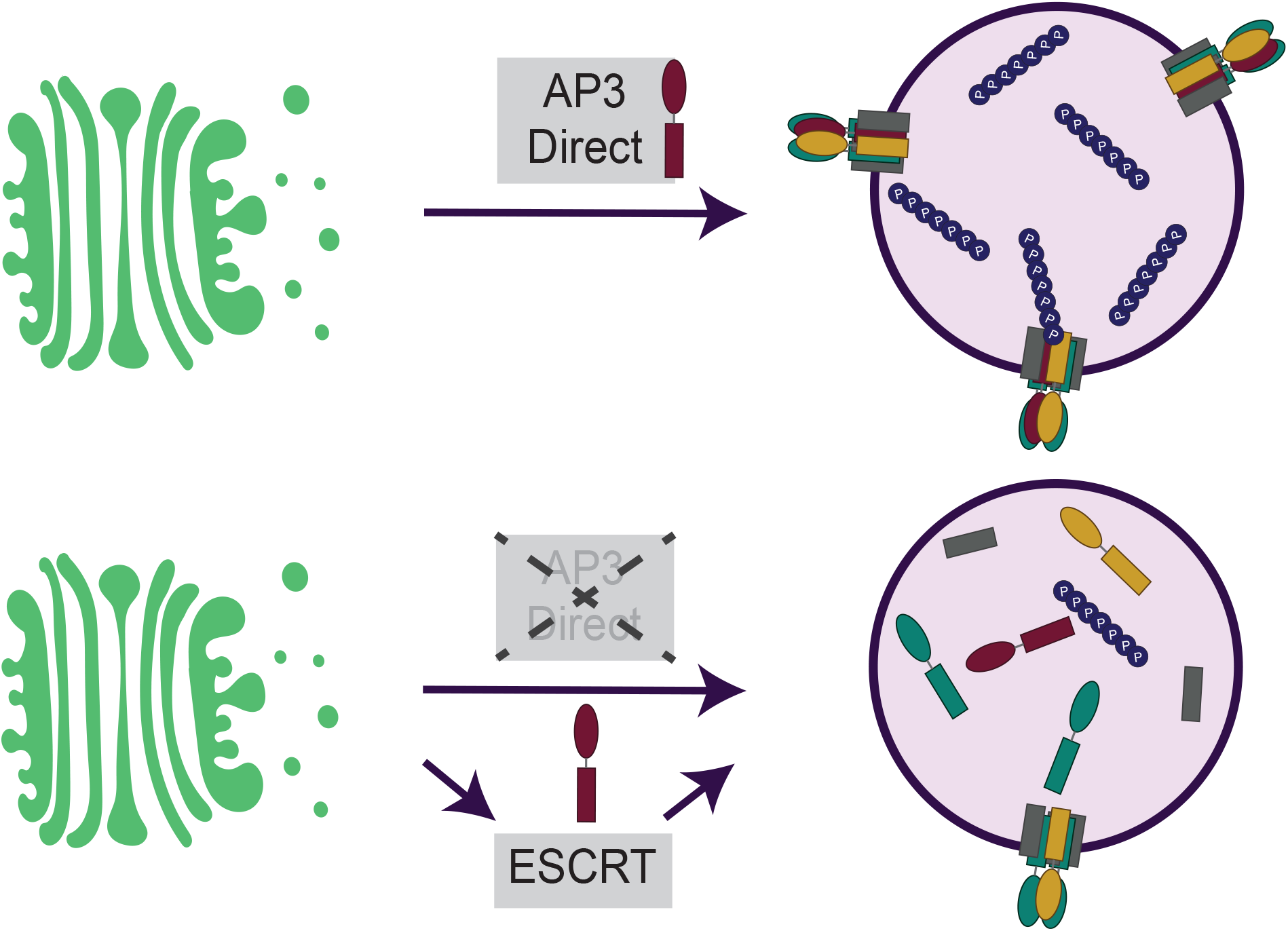
GFP-Vtc5 is localized to the vacuole membrane via the AP-3 complex. In the absence of AP-3, mislocalized GFP-Vtc5 could be localized at the plasma membrane or elsewhere in the cell prior to recognition by the ESCRT complex which delivers it to the vacuole lumen for Pep4-dependent degradation. However, we cannot rule out the possibility that Vtc5 is incorrectly inserted at the vacuole membrane prior to ESCRT-dependent internalization. See text for details.

We found that degradation of Vtc5 in AP-3 mutants requires the Doa4 deubiquitinase. The involvement of Doa4 suggests that mislocalized Vtc5 is subject to regulation by ubiquitylation. In support of this assertion, several large-scale studies have identified ubiquitylation sites in the middle and C-terminal regions of the protein^64^. We speculate that these sites may be targets of the Rsp5 E3 ubiquitin ligase that has been implicated in the ESCRT-dependent delivery of proteins to the vacuole lumen^65-67^. Notably, Vtc5 is also rich in phosphoserines targeted by both the Cdk1 (cell cycle) and Nkk1(nitrogen metabolism)^64^. These modifications could modulate Vtc5 localization in wild-type or AP-3 mutant cells or may instead be involved in regulating its activity towards the core VTC complex.

### Regulation of the core VTC complex

In contrast to GFP-Vtc5, we found that localization of GFP-Vtc3 to the vacuole membrane was not impacted by disruption of AP-3 and GFP-Vtc4 was only partially affected. However, deletion of AP-3 subunits still reduced GFP-Vtc3/4 protein levels and the increased appearance of free GFP in western blots, albeit to a lesser degree than that observed for GFP-Vtc5. We suggest that transport of core VTC subunits occurs through multiple transport routes, although these may function redundantly with AP-3. This type of dual regulation has been documented for other AP-3 cargos such as Sna4^55^ and Ypq1^68^. It is also possible that changes observed in core subunits stem from mislocalized Vtc5. There is strong evidence that maximal polyP production requires VTC localization to the vacuole. However, polyP has also been detected in variable amounts at the plasma membrane, cytoplasm, mitochondria, and nucleus^14^. Whether vacuolar polyP is somehow transported to these compartments or if it is made by VTC residing locally remains an open question. In the latter case, it is possible that Vtc5 association (or lack thereof) with core VTC in these areas also determines subcellular concentrations of polyP outside of the vacuole. In future work it will be intriguing to identify pathways that regulate localization of GFP-Vtc2, which is found at the plasma membrane or peripheral endoplasmic reticulum in phosphate replete conditions, becoming enriched at the vacuole only under conditions of phosphate starvation^9, 12, 15^.

### AP-3 impact on polyP levels

Our work shows that AP-3 mutant cells have reduced levels of polyP. This was true in both a wild-type background and under conditions of Vtc5 overexpression. These findings are consistent with work from Feimoser *et al*. identifying AP-3-encoding genes in a genome-wide screen for deletion mutations with defects in polyP accumulation^60^. Interestingly, deletion of *DOA4* in an AP-3 mutant background rescued both GFP-Vtc5’s localization to the vacuole membrane and restored wild-type levels of full-length GFP-Vtc5 protein without restoring polyP levels. In fact, polyP levels in *apl5*Δ *doa4*Δ double mutants were lower than in either single mutant. While this may seem counterintuitive, we suggest that Vtc5 is not correctly positioned within the vacuole membrane under these circumstances and unable to stimulate VTC activity.

The reduction in polyP levels seen in AP-3 is not as dramatic as that observed in cells lacking Vtc5 altogether, suggesting either residual localization of Vtc5 to the vacuole membrane in AP-3 mutants, or that mislocalized Vtc5 remains competent to stimulate core VTC activity to some degree. Notably, Freimoser *et al*. identified over 200 additional genes that impact polyP metabolism^60^, and some of these could function as direct regulators of Vtc5 or AP-3.

We recently described the Apl5 subunit of AP-3 as a target of lysine polyphoshorylation^46^. However, our analysis of mutant Apl5 that cannot be polyphosphorylated suggests that this modification does not impact GFP-Vtc5 delivery to the vacuole or other AP-3 related phenotypes that we tested. Polyphosphorylation of AP-3 may become important under select stress conditions that remain to be identified. Since polyP has been described as having chaperone activity^4^, another possibility is that polyphosphorylation promotes degradation or refolding of a small fraction of AP-3 that is itself mislocalized to the polyP-rich vacuole lumen.

### Conservation of the AP-3 complex and function

The AP-3 complex is highly conserved in mammalian cells in terms of subunit organization and function. For example, the human homolog of Ypq1, PQLC2, is transported to the vacuole membrane by the AP-3 complex when it is expressed in yeast^68^. Notably, mutations in human AP-3 give rise to Hermansky-Pudlak Syndrome. Relevant here is the observation that molecular phenotypes of Hermansky-Pudlak Syndrome patients include a failure to accumulate polyP in platelet dense granules, a type of lysosome-related organelle conceptually similar to the yeast vacuole. Based on the conserved role of AP-3 subunits in polyP metabolism, we suggest that identification and characterization of AP-3 cargoes in human cells that accumulate high levels of polyP may provide unique insights into the human polyP synthetases, the identity of which remains a critical open question in the field.

## EXPERIMENTAL PROCEDURES

### Yeast strains & handling

Yeast strains were constructed using standard techniques via transformation of PCR products containing selectable markers. For gene deletions, PCR analyses were used to confirm the position of gene deletion and the absence of wild-type gene copy. PCR was also used to confirm the correct genomic location of cassettes used for epitope tagging.

Genotypes for all strains used in this study are listed in **Table S1**. For all experiments performed strains were grown in YPD (2% glucose supplemented with 0.005% Adenine and 0.005% Tryptophan).

### Apl5 K-R Mutagenesis

A construct containing the lysine (K) to arginine (R) mutations within Apl5 PASK cluster was created custom by GenScript and used for the experiments listed in **Supplemental Fig 1 and Supplemental Fig 2**. The construct was integrated with a GFP tag or flag tag at the endogenous *APL5* locus using transformation of overlapping PCR products.

### Electrophoresis & Immunoblotting

Methods for protein extraction were described previously^47^ and are summarized here using similar wording for clarity. A BioSpec bead-beater was used to lyse cell pellets corresponding to 3-6 OD_600_ units in the presence of 100 μL of acid washed beads and 300 μL of 20 % TCA (Sigma-Aldrich T6399). 2 x 3-minute pulses were used. The supernatant was recovered, and cells were washed with 300 µL of 5 % TCA (Sigma-Aldrich T6399). The supernatant from the second wash was combined with the first and this mixture was clarified by centrifugation at 4 °C at 16,000 x *g* for 4 minutes. The supernatant was removed, and the pellet was resuspended in SDS-PAGE sample buffer (see buffer recipes section) supplemented with 1/10 volume of 1.5M Tris-HCl pH 8.8 (Tris Base Fisher BP152-5, Hydrochloric Acid Fisher A144-212) and 1/10 volume 1 M DTT (Bio Basic DB0058). Samples were boiled for 5 minutes before an additional centrifugation at 4 °C at 16,000 x *g* for 4 minutes.

The supernatant was recovered and stored or used immediately for SDS-PAGE or NuPAGE analysis (Thermo Fisher NP0336). Where indicated, BioRad TGX Stain-Free™ FastCast™ 10% acrylamide (BioRad 1610183) was used for quantification with total protein quantified in place of a loading control. BioRad TGX acrylamide gels were transferred to nitrocellulose membrane (BioRad 162-0112) and exposures were obtained using a Bio-Rad ChemiDoc system. Non-TGX 12% SDS-PAGE and NuPAGE gels (Thermo Fisher NP0336) were transferred to PVDF membrane (BioRad 162-0177) and exposures were obtained using autoradiography film (Harvard Apparatus Canada DV-E3018). In all cases, Chemiluminescence Luminata Forte ECL (Fisher Scientific WBLUF0500) was employed for detection. All antibodies used for immunoblotting are described in **Table S2**.

### Spot tests

Cells were diluted to 0.1 OD_600_, grown for 4 hours, then diluted to 0.1 OD_600_ prior to being serially diluted 5-fold in H_2_0. 4 μL of each dilution was spotted on the indicated media prior to incubation at 30 °C for 2-3 days. Plates contained the following chemicals/drugs: 0.028% DMSO (VWR CA97061-250), 0.75 mM NiCl_2_ (Fisher N54-250), and 0.003 µM Rapamycin (Sigma-Aldrich R0395-1MG). Images were taken using a Bio-Rad ChemiDoc.

### Microscopy

Live-cell fluorescence imaging was conducted using a Leica DMI 6000 with a Hamamatsu camera using the Volocity® 4.3.2 imaging program. Briefly, cells were diluted to OD_600_ = 0.2 from overnight cultures in YPD. After 3 hours of growth at 30 °C, FM 4-64 (Thermo Fisher T13320; 1.64 mM stock in DMSO) was added to a final concentration of 1.64 µM for an additional 2 hours. Cells were then washed out in YPD for 30 minutes at room temperature. In instances where there was a difference in protein level expression, exposure times were taken at longer intervals to account for this discrepancy (**Fig. 1**). For microscopy in **Fig. 2**, images were taken on a Zeiss AxioObserver 7 with a Hamamatsu ORCA-Flash LT camera using Zeiss Zen 3.0 Pro Software. All images were taken with oil immersion at 63X. All images were then analyzed in FIJI. Backgrounds were subtracted with a rolling ball radius of 50 pixels and images taken with FM4-64 dye were converted from red to magenta.

### Polyphosphate Extractions

Polyphosphate was extracted from yeast pellets containing 8-12 OD_600_ units using an adapted protocol from Bru *et al* 2016^7, 69^. Cells were resuspended in 400 µL of cold LETS buffer (see buffer recipes section). Subsequently, 600 µL of neutral phenol pH 8 (Sigma-Aldrich P4557) and 150 µL of mH_2_O were added. Samples were vortexed for 20 seconds and heated for 5 minutes at 65°C followed by a 1-minute incubation on ice. 600 µL of chloroform (Sigma-Aldrich 472476) was added, samples were vortexed for 20 seconds and spun down at room temperature for 2 minutes at 13,000 x *g*. The top layer was then transferred to a new tube containing 600 µL of chloroform (Sigma-Aldrich 472476), vortexed for 20 seconds and spun down at room temperature for 2 minutes at 13,000 x *g*. The top layer was transferred to a new tube and 2 uL of RNAse A (10 mg/mL; Thermo Fisher R1253) and 2 µL of DNAse I (10 mg/mL; Thermo Fisher AM2222) were added followed by a 1-hour incubation at 37°C. The mixture was transferred to a pre-chilled tube containing 1 mL 100% ethanol (Commercial Alcohols P006EAAN) and 40 µL of 3M sodium acetate pH 5.3 (Sigma-Aldrich S7899). Samples were left at -20°C overnight and then centrifuged for 20 minutes at 13,000 x *g* at 4°C. The pellet was washed in 500 µL of cold 70% ethanol (Commercial Alcohols P006EAAN), centrifuged for 5 minutes at 13,000 x *g* at 4°C then the supernatant was discarded, and the pellet was dried. The pellet was resuspended in 20-30 µL mH_2_O. Samples were mixed 1:1 with polyP loading dye (see buffer recipes section) and electrophoresed on a 15.8% TBE-Urea Acrylamide Gel at 100V for 1 hour and 45 minutes in 1X TBE buffer (see buffer recipes section). The gel was incubated in fixing solution (see buffer recipes section) with toluidine blue for 15 minutes and then destained in destaining solution. PolyP standards (a gift from T. Shiba) were used to assess polyP chain length.

### Statistical analyses

For statistical analyses performed on western blots in **Fig. 3**, a one-way ANOVA was performed with Tukey posthoc tests at 95% confidence intervals. Error bars, p-values and number of biological replicates (n) are defined in the relevant figure legends.

### Buffer recipes

The following buffer recipes are taken from our previous study (Bentley-DeSousa *et al*., 2018) and are listed here again verbatim for convenience.

#### SDS-PAGE Running Buffer (1X working)

100 mL of 10X 1 litre Stock [30.2 g Tris Base (Fisher BP152-5), 188 g Glycine (Fisher BP381-5), 10 g SDS (Fisher BP166)] and 900 mL ddH_2_O. * Please note SDS-PAGE gels don’t resolve polyP shifts*

#### NuPAGE Running Buffer (1X working)

50 mL of 20X 1 litre Stock [209.2 g MOPS (Sigma M1254), 121.1 g Bis-Tris (Sigma B9754), 20 g SDS (Fisher BP166), 12 g EDTA (Sigma ED2SS)], 5 mL of 1M Sodium Bisulfate (Fisher S654-500) and 950 mL ddH_2_O

#### SDS-PAGE Transfer Buffer (1X working)

100 mL of 10X 1L Stock [30.275 g Tris Base (Fisher BP152-5), 166.175 Glycine (BioBasic GB0235], 200 mL Methanol (Fisher A412P-4) and 700 mL ddH_2_O

#### NuPAGE Transfer Buffer (1X working)

50 mL of 20X 1L Stock [81.6 g Bicine (Sigma B3876), 104.8 g Bis-Tris (Sigma B9754), 6 g EDTA Sigma ED2SS)], 200 mL Methanol (Fisher A412P-4) and 750 mL ddH_2_O

#### SDS-PAGE/NuPAGE Sample Buffer: (3X stock)

800 μL of stock [160 mM Tris-HCl pH 6.8 (Tris base Fisher BP152-5, Hydrochloric acid Fisher A144-212), 6% SDS w/v (Fisher BP166), 30% Glycerol (Fisher BP229-4), 0.004 % Bromophenol Blue (Fisher BP115-25)]. **For TCA preps**, 3X is supplemented with 100 μL 1M DTT (BioBasic DB0058), and 100 μL 1.5M Tris-HCl (pH 8.8) [Tris base (Fisher BP152-5), Hydrochloric acid (Fisher A144-212)

The following buffer recipes are used in Bentley-DeSousa *et al*., 2021.

#### TBE (1X working)

200 mL of a 5X Stock [67.5 g Tris Base (Fisher BP152-5), 34.37 g Boric Acid (Fisher BP168-1), and 25 mL 0.5M EDTA (Sigma-Aldrich 03690)]

#### LETS Buffer (1X working)

100 mM LiCl (Fisher L120-500), 10 mM EDTA (Sigma-Aldrich 03690), 10 mM Tris-HCl Ph 7.4 [(Tris Base Fisher BP152-5, Hydrochloric Acid Fisher A144-212)], 20% SDS (Thermo Fisher AM9820)

#### PolyP Loading Dye (6X)

10 mM Tris-HCl pH 7 [(Tris Base Fisher BP152-5, Hydrochloric acid Fisher A144-212)], 1 mM EDTA (Sigma-Aldrich 03690), 30% Glycerol (Fisher BP229-4), and Bromophenol Blue (Fisher BP115-25)

#### Toluidine Blue Fixing Solution

25% Methanol (Fisher A412P), 5% Glycerol (Fisher BP229-4), 0.05% Toluidine Blue (Sigma-Aldrich T3260)

#### Destaining Solution

25% Methanol (Fisher A412P), 5% Glycerol (Fisher BP229-4)

## ACKNOWLEDGEMENTS

We thank A. Mayer for yeast strains, T. Shiba (Regentiss, Japan) for polyP standards and members of the Downey lab for critical reading of the manuscript. We also acknowledge the Cell Biology and Image Acquisition Core funded by the University of Ottawa, Ottawa, Canada and the Canada Foundation for Innovation for microscopy training and expertise. Work for this project was funded by a Canadian Institutes of Health Research (CIHR) Project Grant (PJT-148722), and by an Early Researcher Award from the Ontario Ministry of Innovation and Research to MD. ABD was funded in part by a graduate scholarship from the Natural Sciences and Engineering Research Council of Canada (NSERC).

## FIGURE LEGENDS

**Supplemental Table 1 – Yeast strains used in this work (Excel File Tab 1)**

**Supplemental Table 2 – Antibodies used in this work (Excel File Tab 2)**

**Supplementary Figure S1:**
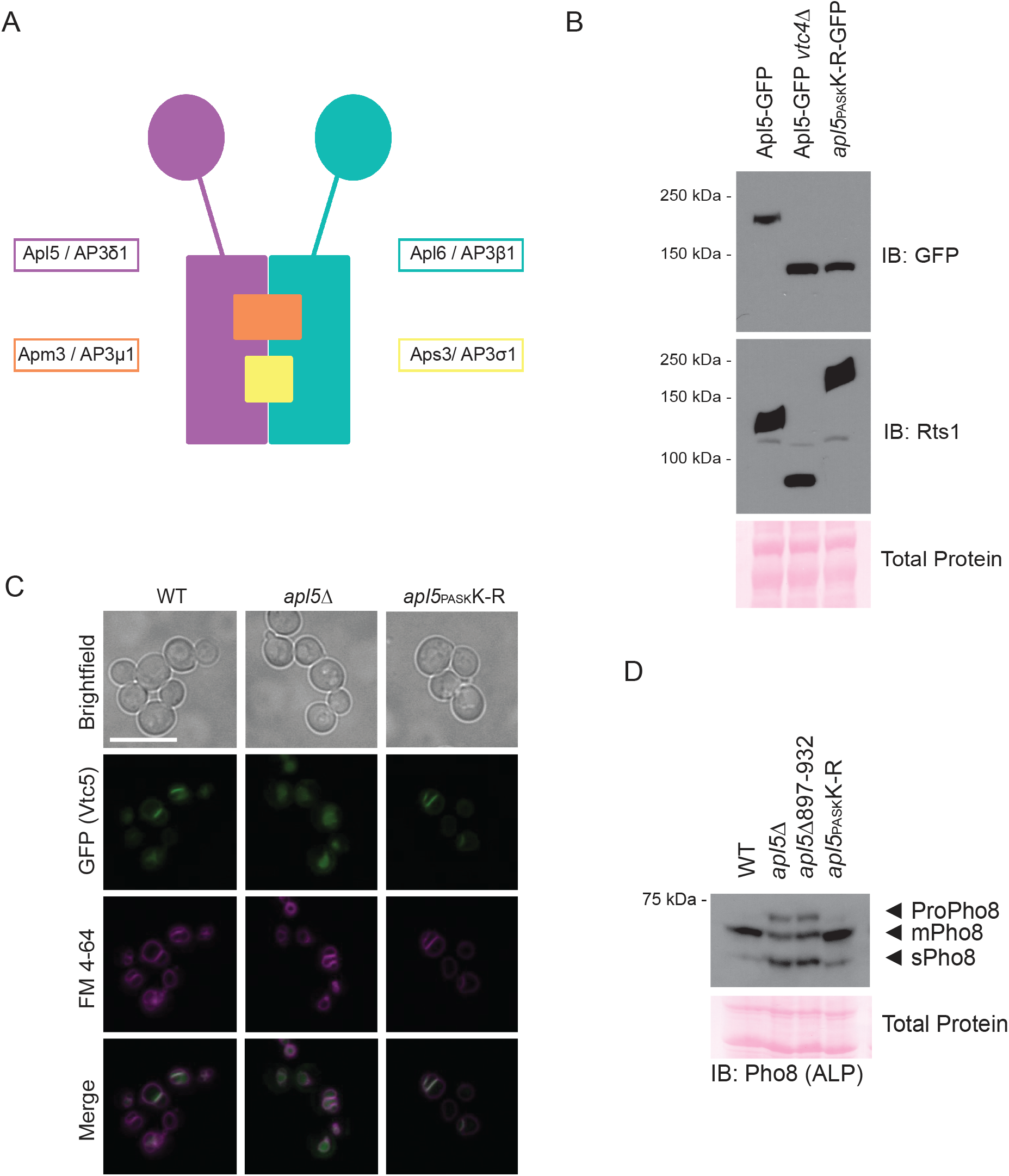
Polyphosphorylation of Apl5 does not impact protein transport. **A)** The AP-3 complex is a conserved heterotetramer which includes two large subunits (Apl5/AP3β1 and Apl6/AP3δ1), one medium subunit (Apm3/AP3μ1), and one small subunit (Aps3/AP3σ1). **B)** Mutating lysines to arginines (K-R) within Apl5’s PASK cluster causes a collapse in the electrophoretic shift on NuPAGE, indicating a loss of polyphosphorylation. Proteins from the indicated strains were extracted using a TCA protein extraction protocol, electrophoresed on a 4-12% NuPAGE gel, and transferred to a PVDF membrane. The membrane was developed with autoradiography film after immunoblotting with an anti-GFP antibody to detect C-terminal Apl5-GFP fusion alleles. Anti-Rts1 serves as a positive control for polyphoshorylation. Ponceau S stain is used as a loading control. **C)** GFP-Vtc5 is localized to the vacuole membrane in Apl5_PASK_K-R mutants. Cells were grown in YPD prior to incubation with FM 4-64, which marks the vacuole membrane, for 2 hours. Cells were then washed with fresh YPD for 30 minutes, transferred to synthetic media, and imaged. Live-cell fluorescence microscopy was performed using a Leica DMI 6000 microscope at 63X with oil immersion. In AP-3 null mutant (*apl5*Δ), green (GFP-Vtc5) images were taken at longer exposure times to account for differences in signal intensity. Images were processed in FIJI. Scale bar represents 10 μm. **D)** Apl5_PASK_K-R does not impact maturation of AP-3 target Pho8. Proteins were extracted from the indicated strains using a TCA protein extraction protocol, electrophoresed on a 12% SDS-PAGE gel, and transferred to a PVDF membrane. The membrane was developed with autoradiography film after immunoblotting with an anti-Pho8 antibody. Ponceau S stain is used as a loading control.

**Supplemental Figure S2:**
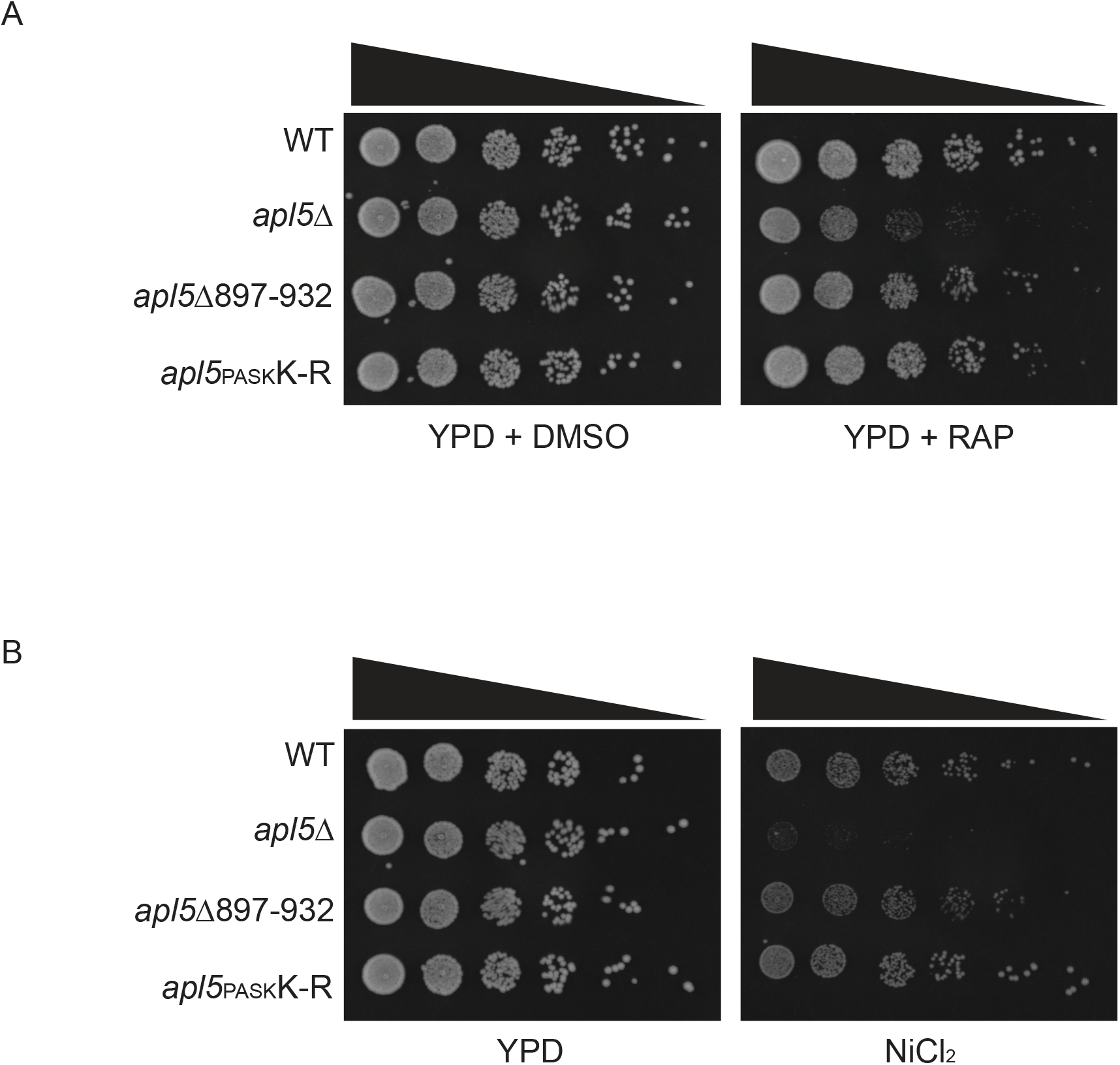
Deletion of the Apl5 PASK cluster does not impact sensitivity to rapamycin or nickel chloride. **A)** Functional AP-3 is required for growth in the presence of 0.003 µM rapamycin. Cells were grown in YPD to log phase and diluted to 0.1 OD_600_ prior to 5-fold serial dilutions in H_2_0. 4 μL of each dilution was spotted on the indicated media prior to incubation at 30 °C for 2-4 days. Images were taken on a Bio-Rad ChemiDoc. DMSO is used as a control for rapamycin treatment. **B)** Functional AP-3 is required for growth in the presence of 0.75 mM NiCl_2_. The same methodology was used as described in Supp. 2A with YPD plates used as a control.

**Supplemental Figure S3:**
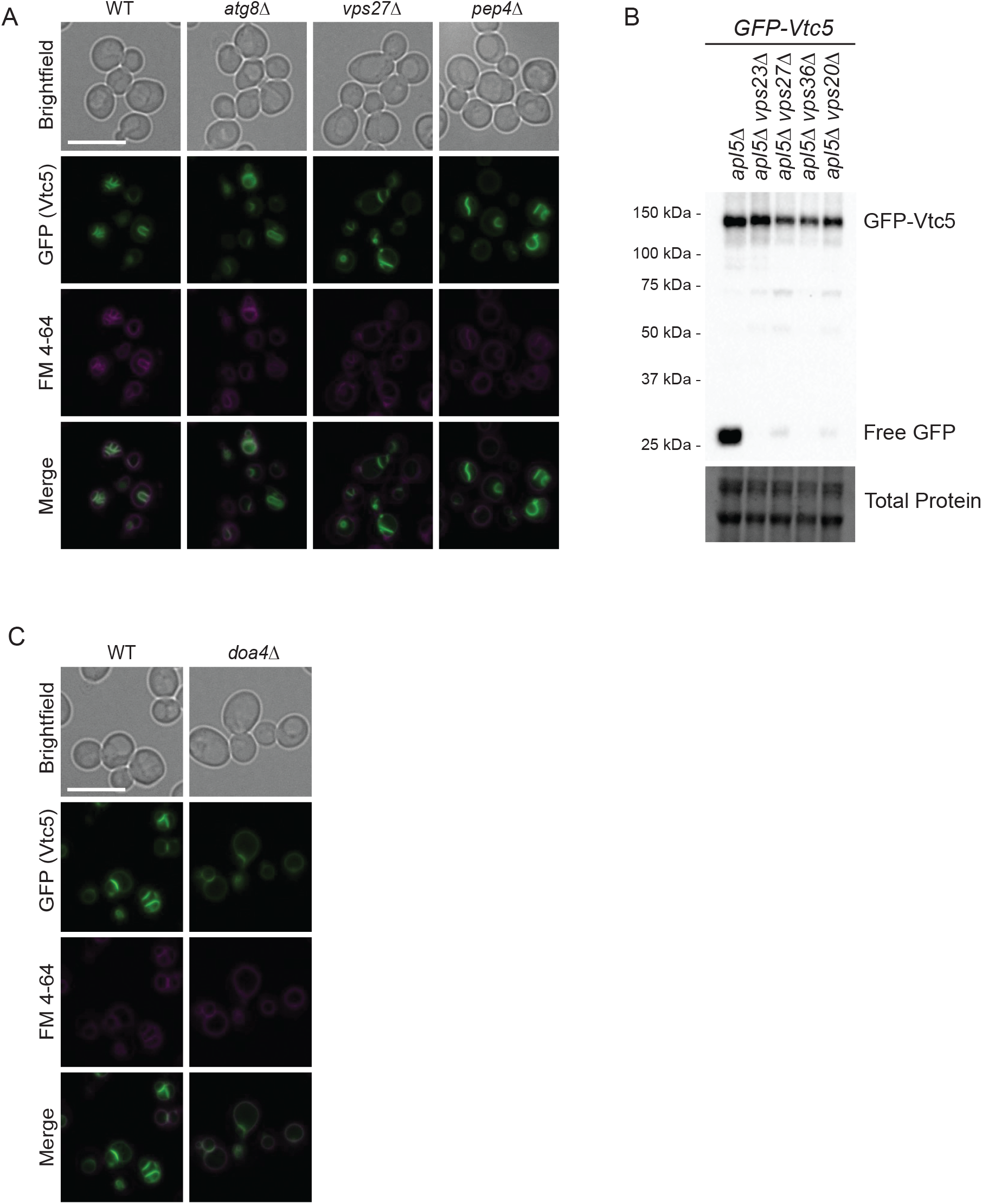
ESCRT complex is responsible for GFP-Vtc5 degradation only when AP-3 is non-functional. **A)** In wild-type cells, GFP-Vtc5 remains localized to the vacuole membrane in autophagy-(*atg8*Δ), ESCRT-(*vps27*Δ), and protease-deficient (*pep4*Δ) cells. The indicated strains were grown in YPD prior to incubation with FM 4-64, which marks the vacuole membrane, for 2 hours. Cells were then washed with fresh YPD for 30 minutes, transferred to synthetic media, and imaged. Live-cell fluorescence microscopy was performed using a Leica DMI 6000 microscope at 63X with oil immersion. Images were processed in FIJI. Scale bar represents 10 μm. **B)** Mutations in ESCRT-0 (*vps27*Δ), ESCRT-I (*vps23*Δ), ESCRT-II (*vps36*Δ), and ESCRT-III (*vps20*Δ) reverse free GFP accumulation in AP-3 mutants expressing GFP-Vtc5. Proteins were extracted using a TCA protein extraction protocol, electrophoresed on a 10% BioRad TGX Stain-Free™ FastCast™ acrylamide gel, and transferred to a nitrocellulose membrane. The membrane was imaged using a BioRad ChemiDoc after immunoblotting with an anti-GFP antibody to detect GFP-Vtc5. Total protein was imaged as a loading control. **C)** Deletion of *DOA4* does not impact wild-type GFP-Vtc5 localization. The same methodology was used as in Supp. 3A.

